# Production of isotopically enriched high molecular weight hyaluronic acid and characterization by solid-state NMR

**DOI:** 10.1101/2023.03.16.532902

**Authors:** Pushpa Rampratap, Alessia Lasorsa, Barbara Perrone, Patrick C.A. van der Wel, Marthe T.C. Walvoort

## Abstract

Hyaluronic acid (HA) is a naturally occurring polysaccharide that is abundant in the extracellular matrix (ECM) of all vertebrate cells. HA-based hydrogels have attracted great interest for biomedical applications due to their high viscoelasticity and biocompatibility. In both ECM and hydrogel applications, high molecular weight (HMW)-HA can absorb a large amount of water to yield matrices with a high level of structural integrity. To understand the molecular underpinnings of structural and functional properties of HA-containing hydrogels, few techniques are available. Nuclear magnetic resonance (NMR) spectroscopy is a powerful tool for such studies, e.g. ^13^C NMR measurements can reveal the structural and dynamical features of (HMW) HA. However, a major obstacle to ^13^C NMR is the low natural abundance of ^13^C, necessitating the generation of HMW-HA that is enriched with ^13^C isotopes. Here we present a convenient method to obtain ^13^C- and ^15^N-enriched HMW-HA in good yield from *Streptococcus equi* subsp*. zooepidemicus*. The labeled HMW-HA has been characterized by solution and magic angle spinning (MAS) solid-state NMR spectroscopy, as well as other methods. These results will open new ways to study the structure and dynamics of HMW-HA-based hydrogels, and interactions of HMW-HA with proteins and other ECM components, using advanced NMR techniques.

## 1. Introduction

Hyaluronic acid (HA) is a linear, negatively charged, non-sulfated glycosaminoglycan (GAG) polysaccharide that contains alternating *N*-acetyl-D-glucosamine (GlcNAc) and glucuronic acid (GlcA) units in the disaccharide repeat →4)-β-D-GlcA*p*-(1→3)-β-D-GlcNAc*p*-(1→ (Figure 1). HA represents one of the major components of the extracellular matrix (ECM) and is found in various tissues, for example navel cords, joint synovial fluid, vitreous humor, dermis and epidermis (Snetkov et al., 2020). HA has high viscoelasticity, and it has the remarkable ability to hold water many times its weight. The molecular weight (MW) of HA is highly variable, ranging from 10^4^ to 10^7^ Da (Fraser et al., 1997; Snetkov et al., 2020). For instance, HA extracted from rooster combs has a MW of 1200 kDa, as compared to 3400-kDa for HA isolated from navel cords. On the other hand, bacterial HA shows a variable MW between 1000 and 4000 kDa (Liu et al., 2011). HA with a MW less than 10 kDa is found in the excreted urine (Buckley et al., 2022).

**Figure 1.**
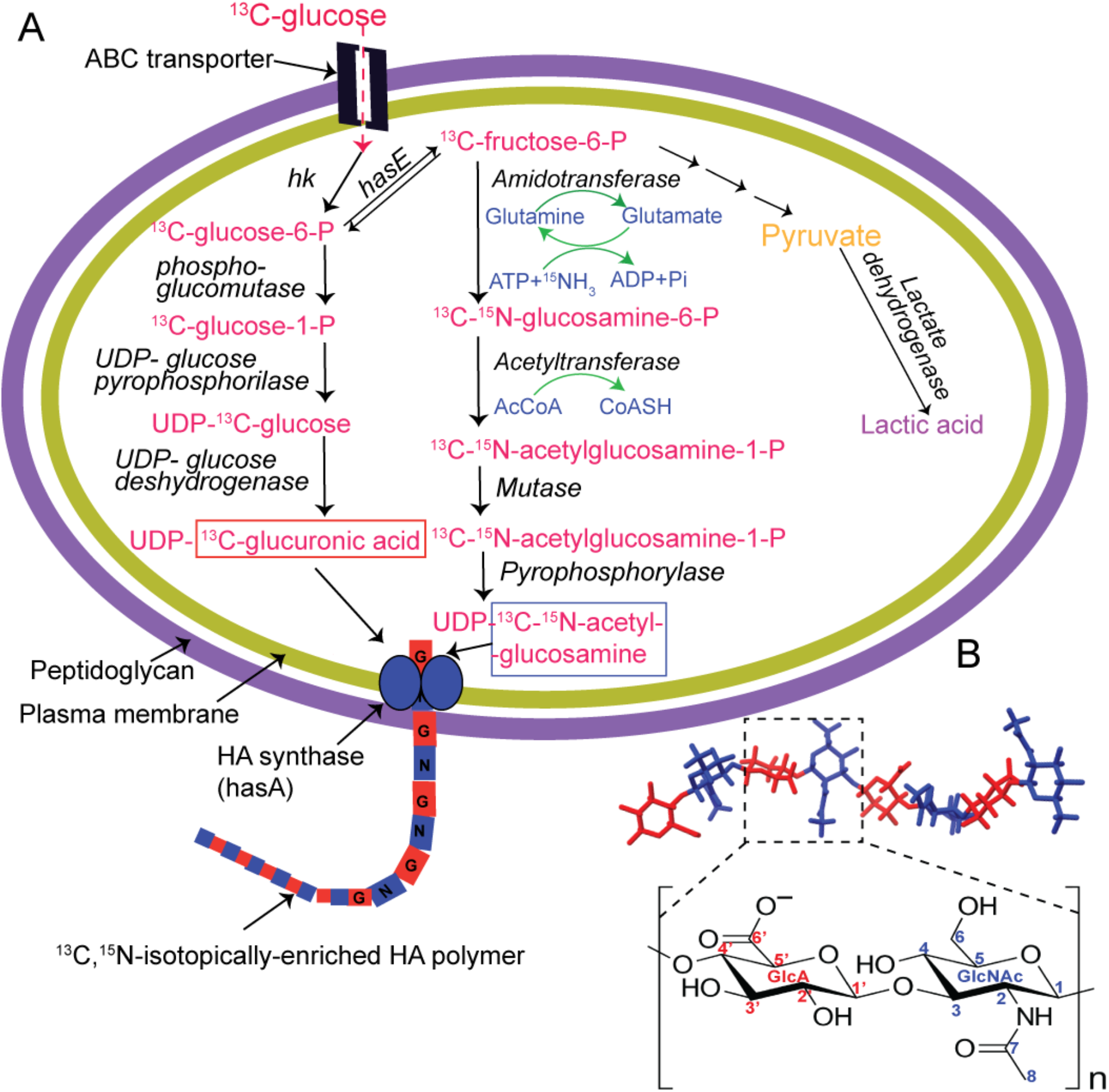
(A) Biosynthetic metabolic pathway for hyaluronic acid in *Streptococcus equi* subsp. *zooepidemicus* and main steps to produce HMW HA. Glucose is first converted into glucose-6-P by hexokinase (hk) which is further used in two different pathways to form UDP-glucuronic acid and UDP-*N*-acetylglucosamine. These sugar nucleotides are then used by the hyaluronic acid synthase (encoded by the *hasA* gene) to produce the HA polymer. Conversion of fructose-6-P to lactic acid is also shown. (B) Molecular structure of HA8 (*top*) from molecular dynamics simulations (PDB 2BVK, (Almond et al., 2006)) and chemical structure of an HA dimer repeat composed of β-1,4-_D_-glucuronic acid (GlcA, red) and β-1,3-*N*-acetyl-_D_-glucosamine (GlcNAc, blue (*bottom*).

The MW of HA has a profound impact on the properties and biological functions of HA. For example, short HA with MW from 0.4 to 4.0 kDa acts as an inducer of heat shock protein expression and has a non-apoptotic function (Xu et al., 2002). HA with MW in the range of 6–20 kDa possesses immunostimulatory and angiogenic activities (Snetkov et al., 2020). HA with MW of 20–200 kDa takes part in biological processes such as embryonic development, wound healing, and ovulation (D’Agostino et al., 2015; Marei et al., 2012), while HA with very high MW (>100 kDa) shows anti-angiogenic activity (Rivera Starr & Engleberg, 2006). Moreover, the mechanisms of interaction of various MW HA with cell surface receptors are an active research area. It has been shown that HA produced by naked mole-rats, which has a very high MW of 6000 kDa, suppresses CD44 signaling, which results in the altered expression of a subset of p53 target genes, suggesting that HMW-HA can act as a cytoprotective molecule (Takasugi et al., 2020).

Thus, it is clear that the MW of HA has a remarkable impact on the observed biological activity. However, the origins of these MW-related effects remain as-yet unclear, limiting our ability to leverage HA as a target for medical interventions. One plausible origin for the different properties of different MWs of HA could be attributed to differences in structural and physicochemical properties. These properties impact the three-dimensional network of the polymer, leading to possible changes in viscosity and viscoelasticity, parameters known to be important in cellular biomechanics (Cowman et al., 2015). Moreover, HA-binding proteins (such as CD44) bind and recognize HA through molecule-specific interactions, which could be modulated by changes in molecular structure in HA of distinct MWs.

Unfortunately, it has remained technically challenging to gain a full understanding of the differences between large- and small-molecular weight HA. Techniques such as X-ray diffraction and solution-state NMR spectroscopy have been used to investigate the molecular structure and conformations of HA (Arnott et al., 1983; Blundell et al., 2005). Molecular dynamics simulations have also been performed to probe the structure and dynamics of HA tetrasaccharides (Donati et al., 2001; Taweechat et al., 2020). However, these reports focus on structure and dynamics of low MW HA, predominantly oligosaccharides. Much less is known about HMW-HA and HA-based hydrogels, mainly due to lack of suitable analysis methods to investigate such semi-solid and complex materials. For instance, HMW-HA is not amenable to crystallization, prohibiting the use of X-ray crystallography. In addition, HMW-HA and polysaccharides in general are too large to be studied by solution-state NMR. Magic angle spinning (MAS) NMR spectroscopy is a unique technique that provides an excellent opportunity to study solid and semi-solid samples, including non-crystalline materials, hydrogels, tissues and whole cells (Murgoci & Duer, 2021; Nestor & Sandström, 2017). Interestingly, MAS NMR analysis has been applied to study HA-based hydrogels and other HA-based (semi)solid samples, including nanoparticles and hydrogels with use in biomedical, cosmetic and pharmaceutical applications (Pouyani et al., 1994; Wu & Liao, 2005). However, these studies employed unlabeled HA, which greatly limits the ability to extract detailed structural information. A major limit of NMR is its inherent low sensitivity for C and N nuclei, making even simple NMR measurements relatively time-consuming. The NMR-active isotopes, including ^13^C and ^15^N, are naturally very low in abundance (1.1 % and 0.4%) (Patching, 2016). Thus, isotopic enrichment with ^13^C and ^15^N is essential to allow application of advanced NMR experiments. While ^13^C and ^15^N isotopic enrichment is a well-established strategy for structure determination of proteins, fewer methods are known to produce other labeled biomacromolecules, including polysaccharides (Pomin et al., 2010; Thomas et al., 2020). Efficient isotope enrichment strategies for HA (and other polysaccharides) promise to facilitate the deployment of similarly advanced NMR methods to this important and growing field of research, spanning biology, pharmaceutical and other (industrial) applications. Some of the impact of such approaches have been illustrated by prior NMR-based studies of ^13^C-enriched polysaccharides in cell walls and other contexts, which have yielded exciting new discoveries regarding their structure and dynamics (Kirui et al., 2022; Pérez García et al., 2012).

Herein we describe our efforts to produce ^13^C- and ^15^N-enriched HMW-HA using Streptococcus bacteria as a host. We hypothesized that by performing bacterial HA production in the presence of ^13^C_6_-labeled glucose and ^15^N-labeled ammonium chloride, a high level of isotope incorporation is achieved, and the resulting HMW-HA is suitable for MAS-NMR analysis. To the best of our knowledge only one protocol is available for the bacterial production of isotopically enriched HA using an engineered *Escherichia coli* K5 strain, which was transfected with HA synthase from *Pasturella multocida* (Blundell et al., 2004). However, HA extraction from this method is laborious and costly due to usage of several organic solvents (DeAngelis et al., 2002) and the yield from the recombinant strains are low as compared to naturally producing strains (Liu et al., 2011).

Interestingly, *Streptococcus equi* subsp*. zooepidemicus* (*S. zooepidemicus*) is a gram-positive bacterium that has extensively been explored for the endogenous production of HMW-HA (Armstrong & Johns, 1997; Fong Chong & Nielsen, 2003; Patil et al., 2011). In contrast to other bacterial systems (Agrobacterium, E. coli and Lactococcus), which need to be genetically modified to express genes that encode for HA biosynthetic pathways, *S. zooepidemicus* naturally produces HA as a component of its extracellular capsule (Kass & Seastone, 1944). This capsule serves for adherence and protection, and molecular mimicry to evade the host’s immune system during its infection process (Wessels et al., 1991). In *S. zooepidemicus*, HA is produced from precursors uridine diphosphate (UDP)-glucuronic acid and UDP-N-acetylglucosamine, which are in turn generated from glucose. Figure 1A details the biosynthesis of HA by Streptococcus bacteria, showing how both GlcA and GlcNAc building blocks originate from glucose (Liu et al., 2011; Ucm et al., 2022). Therefore, we hypothesized that ^13^C_6_-labeled glucose can serve as a starting material for the production of fully ^13^C-labeled HA. Using several enzyme-catalyzed steps, ^13^C-glucose may be converted to ^13^C-labeled UDP-GlcA and UDP-GlcNAc. Because acetyl-CoA also originates from glucose, the ^13^C isotope may also be included in the *N*-acetyl moiety of UDP-GlcNAc. ^15^N isotopes are incorporated through the production of ^15^N-labeled glutamine and subsequent enzymatic transfer by an amidotransferase (Figure 1A). Subsequently, HA synthesis occurs at the plasma membrane, and nascent HA is extruded into the extracellular space (Ghiselli, 2017). Here we report our efforts to explore the above approach, which yielded a practical and efficient method to produce isotopically enriched HMW-HA. Also, we discuss the characterization of the obtained isotopically enriched HMW-HA using solution and solid-state MAS-NMR.

## 2. Materials and methods

### 2.1 Production and purification of ^13^C and ^13^C-^15^N labeled HA

The production protocol was performed in a biosafety level 2 (BSL-2) laboratory. *Streptococcus equi* subsp. *zooepidemicus (S. zooepidemicus)* (DSM 20727) was obtained from Leibniz Institute DSMZ-German Collection of Microorganisms and Cell Cultures. It was cultivated on slants of streptococcus agar containing sheep blood agar. An inoculum was prepared by growing a bacterial culture in 10 mL of 3 % (w/v) tryptone soy broth (TSB) and 1% (w/v) glucose in a 20 mL test tube and incubation at 37 °C for 24 hours. This overnight culture was used to inoculate 100 mL of HA production medium (for ^13^C incorporation: ^13^C-glucose, 25 g/L; casein enzyme hydrolysate, 20 g/L; yeast extract, 5 g/L; NaCl, 2.5 g/L; K_2_HPO_4_, 3 g/L; MgSO_4_·7H_2_O, 5 g/L; pH 7.5; for ^13^C and ^15^N incorporation: ^13^C-glucose, 25 g/L; ^15^NH_4_Cl, 4 g/L, casein enzyme hydrolysate, 14 g/L; yeast extract, 4 g/L; NaCl, 2 g/L; K_2_HPO_4_, 3 g/L; MgSO_4_·7H_2_O, 2 g/L; pH 7.5). The fermentation medium and glucose solution were sterilized by autoclaving and mixed after cooling. The inoculated flask was incubated at 37 °C and 240 rpm shaking for 24 h. After incubation, the broth was diluted with water (1 V) and clarified by centrifugation at 25000 x *g* for 10 min at 4 °C. The supernatant was treated with ethanol (3 V) to precipitate the HA polysaccharides. After filtration using a 0.45 μm syringe filter, the off-white HA precipitate was re-dissolved in 40 mL of MilliQ water and dialyzed in MilliQ water by using a 3.5 kDa cut-off dialysis membrane (SnakeSkin dialysis Tubing). The dialyzed HA was freeze-dried to yield a dry powder.

### 2.2 Molecular weight determination

The MW of HA was determined by gel permeation chromatography (GPC). The analysis was performed on an Agilent Technologies 1200 Series using three PSS (polymer standard service, GmbH) Suprema columns (100, 1000, 3000 Å, 300 × 8 mm × 10 μm), with 40 °C column temperature. The eluate was monitored by a refractive index (RI) detector. The mobile phase was 0.05 M NaNO_3_ at a flow rate of 1 mL/min. 0.1 mg of freeze-dried HA was dissolved in 1 mL of MilliQ water, and samples of 10 μL volume were Injected. Ethylene glycol 0.5% was used as an internal standard. Calibration was performed using a pullulan series (PSS-pulkit-12, Polymer Standard Service, GmbH) with a MW in the range of 1.03–708 kDa. Data analysis was done using PSS winGPC UniChrom software and plotted in Microsoft excel.

### 2.3 HA enzymatic hydrolysis

For the preparation of HA oligosaccharides, 10 mg of ^13^C-HA was dissolved in 2 mL of digestion buffer (150 mM NaCl, 100 mM NaOAc, pH 5.2) and incubated at 37 °C (Mahoney et al., 2001). Ovine testicular hyaluronidase (Sigma-Aldrich, Darmstadt, Germany) was added (0.5 mg; 400-1000 kU in 2ml digestion buffer) and reaction was stopped after 16 hours by boiling the solution for 10 min. Protein precipitate was removed by centrifugation and filtration (0.45 μm syringe filter).

### 2.4 Determination of ^13^C and ^15^N metabolic incorporation by LC-MS analysis

To quantify the ^13^C and ^15^N incorporation in labeled HA, liquid chromatography coupled with mass spectrometry in negative ion mode (LC-MS) was performed (Waters Corporations, Acquity UPLC). Enzymatically hydrolyzed HA oligosaccharides were diluted in MilliQ water to a concentration of 0.1 mg/mL and subjected to LC-MS analysis. An Acquity UPLC BEH Amide 1.7 μm column (2.1 x150mm Column, Waters Corporation) was used in combination with eluents A (0.1% formic acid in H_2_O) and B (0.1% formic acid in acetonitrile), 25 min run (flow rate 0.3 mL/min) with a linear gradient of 10 min from 5% to 50% of B with subsequent increase to 95% B for 3 min. The MS/MS fragmentation experiments were performed using same column as mentioned above. Upon chromatography an eluent was introduced into ESI ion source where the following conditions were set: heater temperature 80°C, capillary temperature 150°C, capillary voltage 2 kV and collision energy ramp was set to 15 (eV) to 50 (eV). The MS/MS spectra were obtained in both positive and negative ion mode in a mass range from 25 to 1000 m/z. The hydrolyzed ^13^C labeled and unlabeled HA oligosaccharides were used for the MS/MS experiments. Data were processed and analyzed by MassLynx software and plotted in MATLAB.

### 2.5 Solution-state NMR spectroscopy

Solution NMR samples were prepared from lyophilized ^13^C-HA powder in 99 % D_2_O (Sigma, Sigma-Aldrich, Darmstadt, Germany): 5 mg of HA was dissolved in 0.5 mL of D_2_O. All spectra were recorded at 25°C on a Bruker Biospin Avance NEO (600 MHz for ^1^H) spectrometer, equipped with a double resonance BBO CryoProbe and using the zgpg30 Bruker pulse program. The experimental parameters used for acquisition were: 90° pulse widths of 12 μs for ^1^H and 10 μs for ^13^C; 1024 number of scans; and 65k number of points. For ^15^N NMR experiments, 8mg ^13^C, ^15^N-enriched HA was dissolved in 0.5 mL of 90% H_2_O, 10% D_2_O. A 2D ^1^H-^15^N HSQC (heteronuclear single quantum coherence) spectrum was recorded using the standard Bruker pulse program (hsqcetgpsi2). The parameters used for acquisition were: 90° pulse widths of 12 μs for ^1^H and 24 μs for ^15^N; 32 number of scans; and 2k and 256 number of points in the direct and indirect dimension respectively. Solution NMR spectra were referenced to DSS and liquid ammonia for ^1^H and ^13^C and, ^15^N respectively. Data were processed and analyzed using Topspin and MestreNova 14.0 software.

### 2.6 Solid-state NMR spectroscopy

Solid-state NMR measurements were performed on an AVANCE NEO 600 MHz (14.1 T) spectrometer from Bruker Biospin, equipped with a 3.2-mm EFree HCN MAS Probe or 4-mm CMP MAS Probe. The corresponding Larmor frequencies were 600.130 and 150.903 MHz for ^1^H and ^13^C, respectively. 8 mg of ^13^C-labeled HA powder were packed in 3.2-mm zirconia rotors whereas 14 mg of the same were packed in 4-mm rotors. Samples were hydrated by adding two-fold excess D_2_O (8 mg of HA dissolved in 16 μL of D_2_O or 14 mg of HA dissolved in 28 μL D_2_O) (99%, Sigma-Aldrich, Darmstadt, Germany), and kept at room temperature for 24 hours prior to solid-state NMR measurements. Additional Kel-F inserts were placed between the sample and the drive cap. ^1^H and ^13^C chemical shifts were referenced to aqueous DSS, and ^15^N referenced to liquid ammonia, using the indirect method, by measuring adamantane ^13^C signals (Harris et al., 2008). ^13^C 1D direct excitation (DE) experiments with high power ^1^H decoupling and ^13^C–^1^H 70– 100 ramped 1D cross-polarization (CP) experiments (with 0.5 ms contact time) were performed using the 3.2-mm EFree HCN MAS Probe, with 50 kHz ^13^C nutation frequency (5 μs 90◦ pulse), 3 s repetition delay, 2048 scans and 0.015 s acquisition time, at 10 kHz spinning frequency and 277 K. During acquisition, proton two-phase pulse-modulated (TPPM) decoupling was around 83 kHz (Bennett et al., 1995; Scholz et al., 2009). The 2D ^13^C-^13^C cross polarization (CP) dipolar-assisted rotational resonance (DARR) solid-state NMR spectrum was acquired using a 4-mm Bruker CMP MAS probe, at 10 kHz spinning frequency and temperature at 298 K. A 70%-100% ramped ^1^H–^13^C CP step with 4 ms contact time and 500 ms DARR ^13^C–^13^C mixing was used, with 4.5 μs 90°carbon pulse and 2.8 μs 90° proton pulse. Frequency-swept TPPM proton decoupling during acquisition was around 88 kHz, repetition delay 3 s and number of scans 256 per datapoint. The acquisition time was 0.040 s in the direct dimension and 0.007 s in the indirect dimension, leading to a truncation of the indirect FID, which manifested as peak broadening in the final spectrum. For the ^15^N ssNMR experiments 12 mg of ^13^C, ^15^N-labeled HMW HA powder was packed into 3.2 mm zirconia rotor. A ^15^N–^1^H 70–100 ramped 1D CP experiment (with 1 ms contact time) was performed using 50 kHz ^15^N nutation frequency (5 μs 90◦ pulse), 4 s repetition delay, 2048 scans and 0.022 s acquisition time, at 10 kHz spinning frequency and 277 K. During acquisition, proton TPPM decoupling was around 83 kHz. 2D ^15^N–^13^C TEDOR spectra were recorded using a 5 μs 90° carbon pulse, 2.5 μs 90° proton pulse, 5 μs 90° nitrogen pulse, 2 ms contact time and TEDOR block total durations of 1.6 and 2.4 ms (Jaroniec et al., 2002). Every TEDOR 2D spectrum was recorded with 64 scans and 0.010 (1170 points) and 0.005 s (130 points) acquisition time in the direct and indirect dimension, respectively. TPPM proton decoupling during <u>TEDOR,</u> and acquisition was around 100 kHz. Spectra were processed with Bruker Topspin and NMRPipe software (Delaglio et al., 1995). NMR spectral analysis and peak assignment were performed with CcpNmr version 2.4 (Stevens et al., 2011).

## 3. Results and discussion

### 3.1 Production of isotopically enriched HA with high molecular weights

We set out to investigate the production of HMW-HA using *S. zooepidemicus* using unlabeled glucose. Standard growth conditions for *S. zooepidemicus* include incubating the culture for 24 h at 37°C and at neutral pH 7.5, in the presence of regular D-glucose. Firstly, we investigated the impact of increasing the fermentation time to 48h to see if longer periods could affect HA production and yield (Table 1, entries 1 and 2). We observed that the HA yield significantly dropped by more than 50% from ∼105 mg to ∼35 mg upon increasing fermentation time (per 100 mL fermentation broth). As reported (Rivera Starr & Engleberg, 2006; Samadi et al., 2022), when fermentation media become deprived of the nutrients, bacterial cells can utilize HA as a carbon source leading to a decrease in HA concentration. Also, during the bacterial production of HA, a hyaluronidase is released from the cells into the culture broth, which may be responsible for enzymatic degradation of HMW-HA upon longer fermentation times (Mausolf et al., 1990). These shorter HA fragments may be removed during the purification steps, resulting in a diminished yield of HMW-HA.

**Table 1.** Overview of the HA samples that were isolated from different growth conditions.

| Batch | Fermentation duration (hours) | Aeration (Culture filling volume) * | Yield (mg/100ml) | Molecular weight distribution (Dalton) | Contamination |
| --- | --- | --- | --- | --- | --- |
| **HA** | 1-2 % seed culture |  |  |  |  |
| 1 | 24 | 1/5 | $105 \pm 10^{\S}$ | NA | NA |
| 2 | 48 | 1/5 | $35 \pm 05^{\#}$ | $1 \times 10^4 - 1 \times 10^6$ | NA |
| 3 | 24 | 1/2 | $65 \pm 10^{\S}$ | $1 \times 10^4 - 5 \times 10^5$ | NA |
| **$^{13}\text{C}$-HA** | 1-2 % seed culture |  |  |  |  |
| 4 | 24 | 1/5 | $95 \pm 10^{\S}$ | $1 \times 10^4 - 1.5 \times 10^6$ | - |
| 5 | 24 | 1/2 | $55 \pm 05^{\#}$ | $1 \times 10^4 - 1 \times 10^6$ | +++ |
| **$^{13}\text{C}$-HA** | 5 % seed culture |  |  |  |  |
| 6 | 24 | 1/5 | $70 \pm 05^{\#}$ | $1 \times 10^4 - 1.2 \times 10^6$ | ++ |
| 7 | 24 | 1/2 | $50 \pm 05^{\#}$ | $1 \times 10^3 - 4 \times 10^4$ | ++++ |
| **$^{13}\text{C}$, $^{15}\text{N}$-HA** | 1-2 % seed culture |  |  |  |  |
| 8 | 24 | 1/5 | $50 \pm 05^{\#}$ | NA | - |

$\S$ Average of three different batches. $\#$ Average of two batches. \* Culture filling volume: 1/5 (i.e., 500 ml Erlenmeyer flask containing ~100 mL of culture) and 1/2 (i.e., 250 mL Erlenmeyer flask containing ~ 100 mL of culture)

Aeration and agitation during streptococcal fermentation are also found to impact HA production (Liu et al., 2011). To investigate these in our system, we performed fermentation experiments in flasks of different volumes. Compared to a low aeration (i.e., 250 mL Erlenmeyer flask was used for 100 ml culture, batch 3), the high aeration culture (i.e., 500 mL Erlenmeyer flask was used for 100 ml culture, batch 1) gave higher HA yield. This was consistent, as the cultures with low aeration (1/2 filling volume) repeatedly gave lower HA yields also in the follow-up experiments (entries 5 and 7, Table 1).

Having established suitable growth conditions for HA production, we set out to investigate the ^13^C-, ^15^N-, and ^13^C,^15^N-enrichment of HMW-HA using *S. zooepidemicus*. Fully ^13^C-labeled glucose (containing 6 ^13^C atoms) was added to the cell culture medium, and fermentation was carried out with different amounts of starting (seed) culture (from 1% to 5%) to investigate the impact of culture dilution. Interestingly, when using a larger amount of seed culture, a slight decrease in HA yield was observed, from ∼95 mg to ∼70 mg per 100 mL (entries 4 and 6, Table 1). In addition, we observed an increase in the production of a contaminant (see below). Especially under conditions of low aeration, an increased concentration of this contaminant was obtained (entries 5 and 7, Table 1). As discussed below, we postulate that this contaminant is lactic acid, which is a well-known by-product of HA fermentation (Liu et al., 2011). If the fermentation conditions are not controlled (i.e., pH, aeration and duration), most of the carbon sources are utilized to produce lactic acid (Lu et al., 2016). In fact, by doing further optimizations we found that limiting the amount of seed culture to 1-2% and increasing aeration was the best condition to obtain good yields and suppress the presence of the contaminant (batch 4, Table 1). Finally, these optimized fermentation conditions were also used to perform a double-labeling experiment, where both ^13^C-glucose and ^15^N-ammonia were added. These conditions resulted in the isolation of isotope-enriched HA in decent yields (∼50 mg per 100 mL).

We determined the MW distribution of all HA polysaccharide samples using gel permeation chromatography (GPC) (Table 1, Figure S1). It can be clearly seen that ^13^C-labeled HA contains a mixture of several polymeric populations in a MW range from 10 kDa to more than 1 MDa. Gratifyingly, the addition of ^13^C-glucose did not affect the molecular weight distribution of the isolated HA. In contrast, we found that aeration and agitation have an impact on the MW, as increased aeration produced HA with more than 1 MDa MW, whereas under decreasing aeration conditions the majority of HA was below 100 kDa (Figure S1).

### 3.2 Characterization of isotopically enriched HA by LC-MS and solution-state NMR

To characterize the produced HA samples, and to determine the efficiency of ^13^C and ^15^N metabolic incorporation, we performed liquid chromatography mass spectrometry (LC-MS). First the polysaccharides were hydrolyzed with a hyaluronidase enzyme that specifically cleaves polymeric HA into tetrasaccharides with GlcNAc at the reducing end (Mahoney et al., 2001). Upon digestion for 16 hours, the HA tetramers were found to be the major species. Both unlabeled and labeled samples were digested under these conditions, and negative-ion mass data were collected (Table 2 and S1, Figure 2). For unlabeled HA, the mass was found to be 774.9 Da in accordance with the calculated value of HA tetrasaccharide (m/z 774.9, Figure 2A). For ^13^C-HA, a mass corresponding to a HA tetrasaccharide with twenty-eight incorporated ^13^C atoms (fully labeled) was found as the major peak (52 % relative abundance, m/z 802.9). Additionally, 22 % and 16 % of HA tetrasaccharides showed partial ^13^C incorporation, with twenty-six and twenty-seven incorporated ^13^C atoms respectively (m/z 800.9 and 801.9). We hypothesized that this incomplete ^13^C-enrichment is due to varying levels of ^13^C incorporation in the acetyl group in the GlcNAc residue. During the biosynthesis of UDP-*N*-acetylglucosamine from glucose 6-P, an unlabeled acetyl group, derived from unlabeled carbon sources present in the medium, such as the yeast extract, may have been transferred by the acetyltransferase enzyme (Figure 1A). We performed MS/MS fragmentation studies which revealed the near-complete enrichment of both the GlcA and GlcNAc units (Figure S5 and S6). Looking at the individual GlcA and GlcNAc moieties, they are mostly fully enriched (199.1 and 212.2 m/z). The main other species lack just a single ^13^C (m/z 198.1 and 211.2), reflecting either 12% or 23% of the main peak, for GlcA and GlcNAc, respectively. Thus, this analysis shows that the incomplete labeling mostly affects the GlcNAc moiety. However, the fragmentation within one moiety was not observed, such that the exact labeling patterns remains uncertain, in absence of further studies.

**Figure 2.**
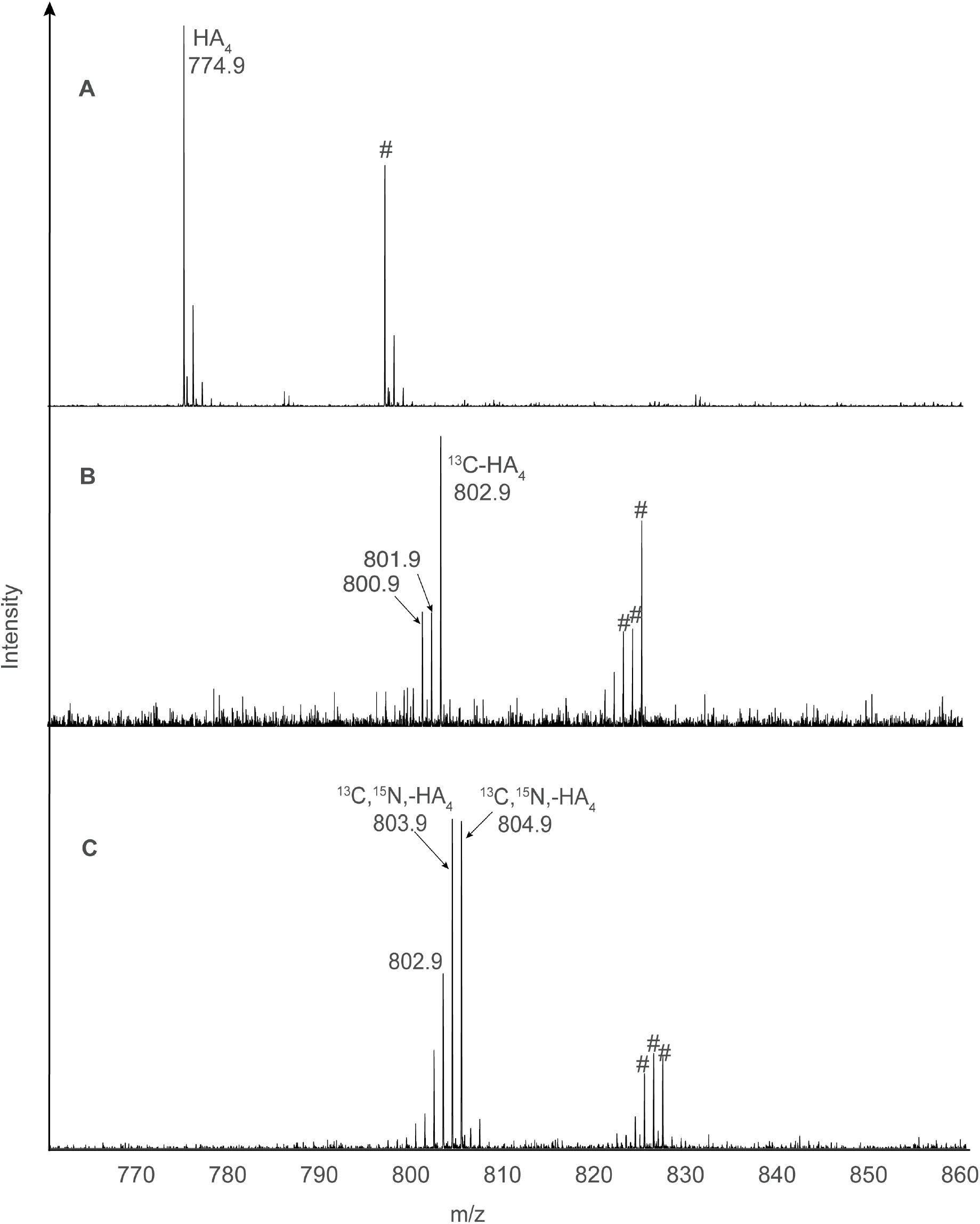
ESI-MS spectra of tetrasaccharide fragments derived from unlabeled HA (panel A), ^13^C-labeled HA (panel B), and ^13^C,^15^N-labeled HA (panel C) showing the monoisotopic mass of their respective monovalent anions measured in negative ion mode. Peaks labeled with hash symbol (#) correspond to the respective monosodium adducts.

**Table 2.** Overview of observed and theoretical molecular masses of unlabeled and isotopically labeled HA_4_ – tetrasaccharides, measured in negative ion mode.

| Digested HA | Observed masses ($\pm 0.2$ Da) | Theoretical mass (Da) |
| --- | --- | --- |
| HA<sub>4</sub> | 774.9 | 775.2 (M-H)<sup>-</sup> |
| $^{13}\text{C}$-HA<sub>4</sub> | 802.9, 801.9, 800.9 | 803.3 (M-H)<sup>-</sup> |
| $^{15}\text{N},^{13}\text{C}$-HA<sub>4</sub> | 804.9, 803.9, 802.9, 801.9 | 805.3 (M-H)<sup>-</sup> |

For the double-labeled ^13^C,^15^N-HA, a mass corresponding to the HA tetrasaccharide with twenty-eight ^13^C atoms and two ^15^N nitrogen atoms (i.e., fully ^13^C, ^15^N-labeled) was found as one of the major peaks (35% relative abundance, m/z 804.9, Figure 2C, Table S1). In addition, three other peaks (34 %, 14% and 10% relative abundance, with m/z values of 803.9, 802.9 and 801.9 respectively, Figure2C, Table S1) were observed which arise from combinations of partial ^15^N and ^13^C incorporation. A complete mass analysis of ^13^C and ^13^C,^15^N incorporation in HA tetrasaccharides is reported in Table S1.

For an additional verification of HA identity and ^13^C enrichment, ^1^H and ^13^C solution-state NMR spectroscopy was performed of intact ^13^C-enriched HMW-HA (Figure 3). The obtained ^13^C-NMR spectrum exclusively contained signals matching those previously reported for HA, and all peaks were assigned (Blundell et al., 2004, 2006; Scott & Heatley, 1999). The signal intensities are relatively low, with the shown spectrum requiring 1000 accumulations. This is due to the slow tumbling rate of the HMW polysaccharides and their lower solubility in water. Whereas this signal loss is in part compensated by the ^13^C enrichment, it is also likely that these solution NMR signals are biased toward HA molecules with lower molecular weights, which constitute a subset of molecules in the sample (Table 1). It is worth noting that ^13^C-labeling in itself is also expected to broaden the lines in these ^13^C NMR spectra (compared to natural abundance samples) due to the presence of ^13^C-^13^C couplings. A ^1^H,^15^N HSQC spectrum was recorded for the ^13^C,^15^N-labeled HA (batch 8), yielding a clear cross peak between the amide nitrogen and proton of the GlcNAc moiety (Figure S4).

**Figure 3.**
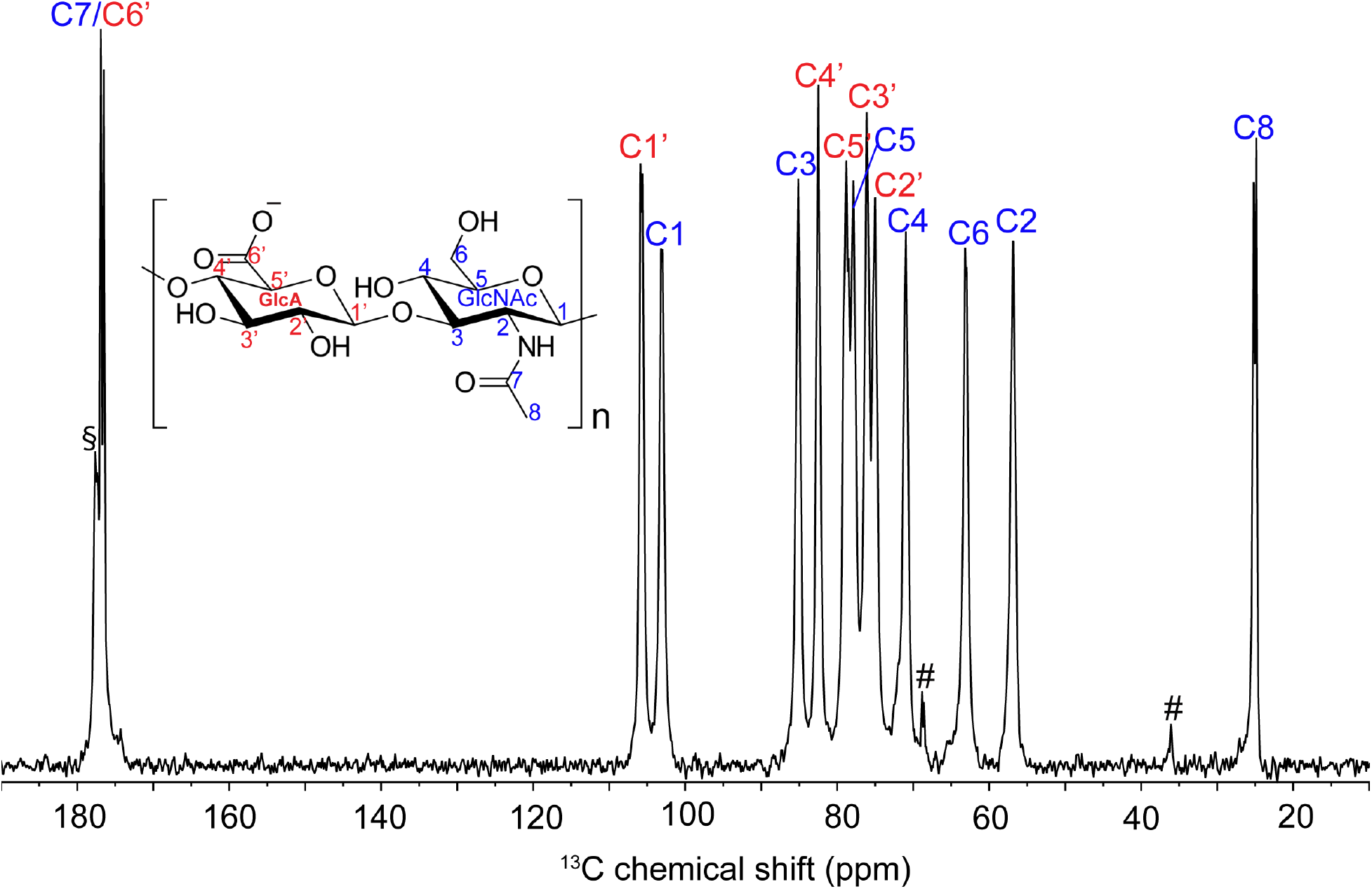
Solution-state ^13^C NMR spectrum of HA (10 mg/mL) along with chemical shift assignment, recorded in D2O at 600 MHz with ^1^H decoupling. The peaks labeled as # are from impurities in the sample, while the peak labeled as § may result from partial labelling. From a peak integration analysis, this second carbonyl peak appears 60% labeled. Spectrum recorded on batch 6 from Table 1.

### 3.3 Characterization of isotopically enriched HMW-HA by MAS NMR

As a complement to solution NMR, MAS NMR is a preferred tool for the analysis of intact HMW polysaccharides. By spinning the sample at the magic angle, one can sidestep the reliance on rapid tumbling of molecules in solution, enabling high-resolution NMR studies of (semi)solids. It can be applied to dry HA powder (as illustrated in Figure S7), but for improved spectral resolution and mimicking the physiological state, hydration of the polysaccharide sample is beneficial (Mandal et al., 2017; Tang et al., 1999). Deuterium oxide was used to hydrate the isotopically enriched HA powder, to attain a two-fold excess of water:HA (2:1, w/w). Figure 4 shows the 1D ^13^C NMR spectra obtained under MAS, recorded with two different types of MAS NMR experiments, i.e., cross polarization (CP) and direct excitation (DE). The observed HA peaks are marked in panel A of the figure, based on solution NMR assignments and verification with 2D ssNMR analysis (see below). It is worth noting that the 1D CP spectrum exhibits lower intensity compared to the DE spectrum. This is primarily because the sample is hydrated, which increases its flexibility and makes it less ideal for CP-based experiments. This conclusion stems from the fact that CP experiments (unlike direct ^13^C excitation) generally only produce detectable signals from relatively rigid molecules (Matlahov & van der Wel, 2018). In the DE ssNMR experiment, an extra peak at around 68 ppm (labeled with an asterisk *) was detected in the initial batch of ^13^C HMW HA that could not be assigned to HA. We hypothesized that this extra peak is likely corresponding to carbon 2 (C2) of lactic acid (LA), a common byproduct during HA fermentation, as discussed above (Figure 1A) (Liu et al., 2011; Lu et al., 2016). The fact that this peak at 68 ppm is present in the DE spectrum, but not in the CP-based spectrum, is an indication that it corresponds to a highly dynamic molecule, which suggests it is dissolved in the aqueous solvent phase. As described above, modification of the fermentation conditions yielded much purer HA samples where this peak almost disappeared (green spectrum in Figure 4B).

**Figure 4.**
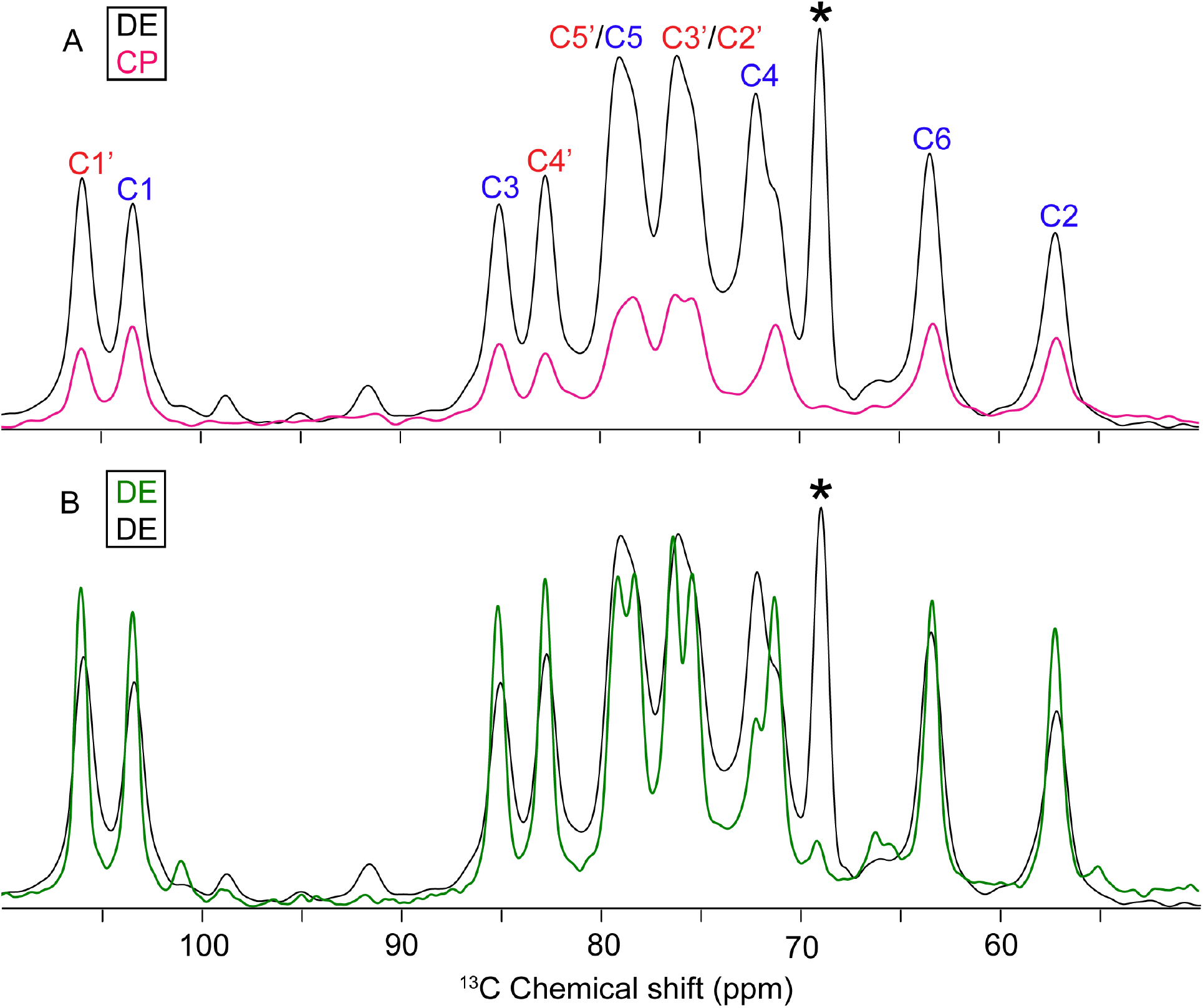
^13^C MAS NMR spectra of hydrated ^13^C-enriched HMW HA. (A) Overlaid cross-polarization (CP, pink) and direct excitation (DE, black) spectra of ^13^C HA sample containing contamination (batch 7 from Table 1). (B) Overlay of the ^13^C DE MAS NMR spectra of batches 7 (black) and 4 (green) of ^13^C-enriched HA, illustrating the reduction in the contaminant peak at 68 ppm (labeled with asterisk; *) in 4. The full DE NMR spectra are included in Figure S2 of the Supplementary Information.

To complement these 1D spectra, we also recorded a CP-based 2D MAS NMR spectrum (batch 4, calculated average MW ∼275 kDa, Figure S1). This 2D ^13^C-^13^C spectrum employed dipolar assisted rotational resonance (DARR) (Takegoshi et al., 2001) recoupling to predominantly detect one-bond ^13^C-^13^C correlations (Figure 5). The 2D MAS NMR spectrum allowed for the verification of the peak assignments, based on the observed pattern of connectivity. For instance, using the 2D spectrum all atoms from the two monosaccharides were identified, highlighting its usefulness in assigning monosaccharide character. The obtained ^13^C chemical shift assignments for both solution- and solid-state NMR of HA are reported in Table 3. Small to modest differences in chemical shift are observed, with maximum differences located in the GlcNAc moiety and especially its exocyclic C7 carbon. It is worth noting that especially such 2D ^13^C-^13^C NMR spectra depend on the ^13^C enrichment of the HMW HA, as accomplished in this work. Mainly one-bond intra-residue correlation peaks can be observed, except for inter-residue cross peaks attributed to the C1-C4’ glycosidic bond (Figure 5 and S3). Several low-intensity multiple-bond intra-residue peaks have also been detected. These peaks mainly belong to the GlcA moiety, indicating that this moiety presents decreased dynamics. Correlations with the carbonyl groups, C5’-C6’ and C7-C8, are shown in Figure S3.

**Figure 5.**
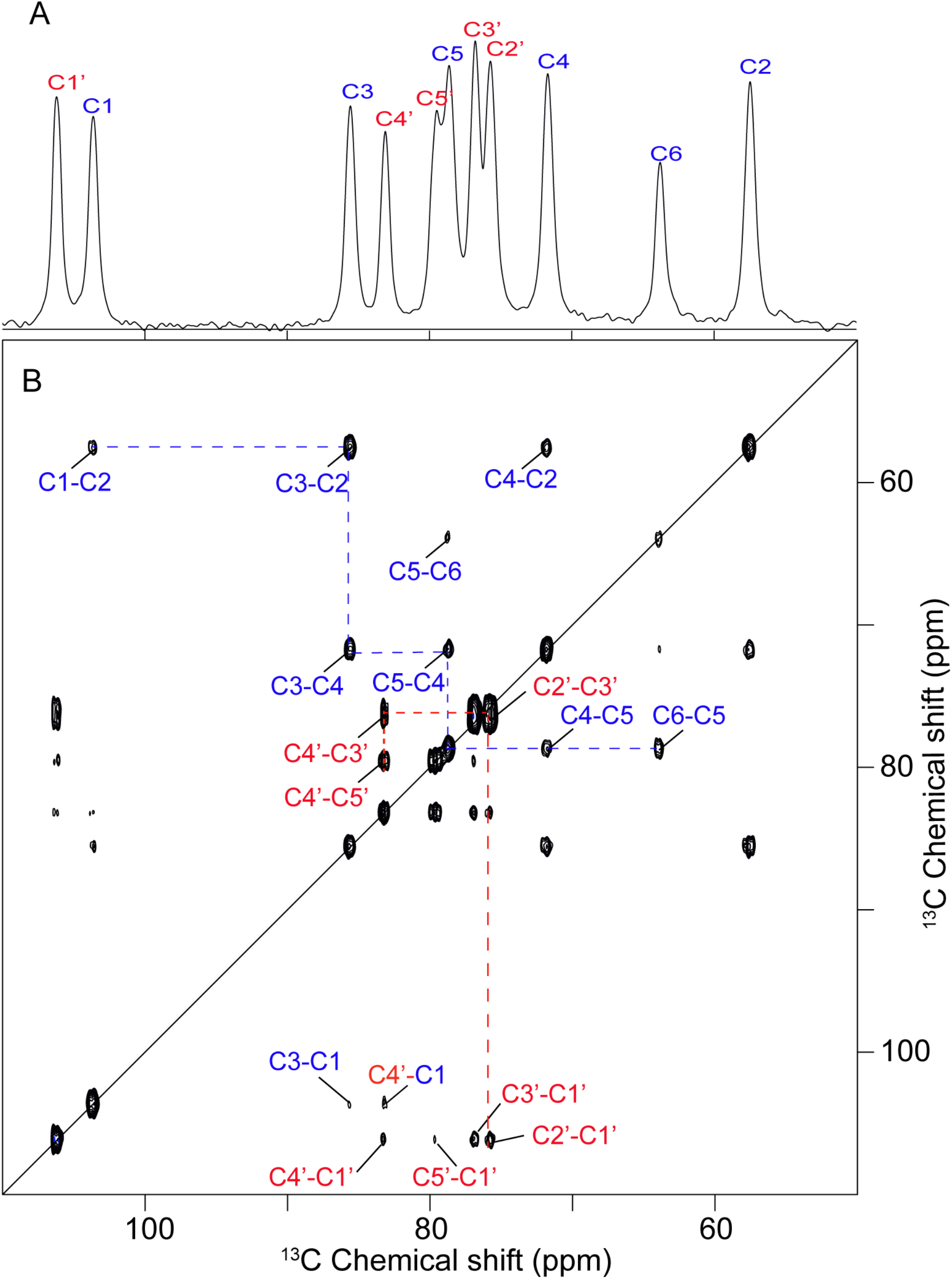
1D and 2D MAS ^13^C and ^13^C-^13^C NMR spectra of ^13^C HMWHA with resonance assignment. (A) 1D CP-MAS NMR spectrum of hydrated HA and (B) 2D DARR spectrum, along with the indicated carbon resonance assignment. Blue labels indicate carbons belonging to GlcNAc and red ones to GlcA. For illustrative purposes, dashed lines show selected carbon connectivities within a single moiety. Spectra recorded on batch 4 from Table 1.

**Table 3.** ^13^C chemical shift assignment of HA (based on solution NMR and CP-DARR ssNMR experiments on batch 6 and 4, respectively, from Table 1).

| Nuclei | $\delta$ (ppm) -<br>CP-DARR | $\delta$ (ppm) -<br>Solution<br>NMR | $\Delta\delta$ (ppm) | Nuclei | $\delta$ (ppm) -<br>CP-DARR | $\delta$ (ppm) -<br>Solution<br>NMR | $\Delta\delta$ (ppm) |
| --- | --- | --- | --- | --- | --- | --- | --- |
| **GlcA** |  |  |  | **GlcNAc** |  |  |  |
| C-1' | 105.8 | 105.7 | 0.1 | C-1 | 103.4 | 103.2 | 0.2 |
| C-2' | 75.5 | 75.1 | 0.4 | C-2 | 57.4 | 57.0 | 0.4 |
| C-3' | 76.6 | 76.2 | 0.4 | C-3 | 85.4 | 85.2 | 0.2 |
| C-4' | 83.0 | 82.5 | 0.5 | C-4 | 71.5 | 71.1 | 0.4 |
| C-5' | 79.3 | 79.0 | 0.3 | C-5 | 78.5 | 78.0 | 0.5 |
| C-6' | 177.2 | 177.0 | 0.2 | C-6 | 63.6 | 63.1 | 0.5 |
|  |  |  |  | C-7 | 177.9 | 177.2 | 0.7 |
|  |  |  |  | C-8 | 25.5 | 25.2 | 0.3 |

To assign the amide group in the GlcNAc moiety we measured a 1D ^15^N CP NMR spectrum on a sample of ^15^N,^13^C-enriched HMW-HA (Figure S7). This spectrum shows one peak around 122 ppm representative of the N-acetyl group. 2D ^15^N-^13^C TEDOR NMR spectra (Jaroniec et al., 2002) with two different mixing times were recorded on the same sample. These spectra show correlations of the nitrogen atom with the carbonyl group (C7) and CH group (C2). At longer mixing time, a correlation with the methyl group (C8) is also visible.

## 4. Conclusion

In summary, we have developed a straightforward method to produce isotopically enriched HMW-HA with a good yield and characterized it by 1D and 2D solution and solid-state NMR. We obtained isotopically enriched HA with a MW in the range of kDa to MDa, which is relevant to the key roles of HA in biology (e.g., as an ECM component) but also for its biomedical and industrial applications. Additionally, by carefully controlling the amount of seed culture and aeration, we were able to significantly limit the presence of a contaminant that was detected by NMR. The use of fully ^13^C-labeled HMW-HA in 2D solid-state NMR was illustrated, which will enable the investigation of the structural properties of HMW-HA without the need for degradation to improve spectral quality. We foresee that this approach can be extended to both HA hydrogels and ECM preparations representing important biological contexts. The employment of isotopic labeling in HMW-HA will facilitate more advanced multidimensional solid-state NMR experiments that can detect and measure HMW-HA in relevant physiological and pathological conditions, as well as in various contexts such as in interaction with HA receptors or other ECM components.

## Acknowledgment

We thank Willem Woudstra (Dept. of Biomedical Engineering at UMCG) for assistance during sample production, Renze Sneep (Stratingh, UG) for helping with LC-MS data acquisition and Leon Rohrbach (ENTEG, UG) for helping with GPC data acquisition. This work was supported by financial support from the Zernike Institute for Advanced Materials at the University of Groningen, including funding from the Bonus Incentive Scheme of the Dutch Ministry for Education, Culture and Science (OCW).

## References

Almond, A., DeAngelis, P. L., & Blundell, C. D. (2006). Hyaluronan: The Local Solution Conformation Determined by NMR and Computer Modeling is Close to a Contracted Left-handed 4-Fold Helix. Journal of Molecular Biology, 358(5), 1256–1269. https://doi.org/10.1016/j.jmb.2006.02.077

Armstrong, D. C., & Johns, M. R. (1997). Culture Conditions Affect the Molecular Weight Properties of Hyaluronic Acid Produced by Streptococcus zooepidemicus. Applied and Environmental Microbiology, 63(7), 2759–2764. https://doi.org/10.1128/aem.63.7.2759-2764.1997

Arnott, S., Mitra, A. K., & Raghunathan, S. (1983). Hyaluronic acid double helix. Journal of Molecular Biology, 169(4), 861–872. https://doi.org/10.1016/S0022-2836(83)80140-5

Bennett, A. E., Rienstra, C. M., Auger, M., Lakshmi, K. V., & Griffin, R. G. (1995). Heteronuclear decoupling in rotating solids. The Journal of Chemical Physics, 103(16), 6951–6958. https://doi.org/10.1063/1.470372

Blundell, C. D., Almond, A., Mahoney, D. J., DeAngelis, P. L., Campbell, I. D., & Day, A. J. (2005). Towards a Structure for a TSG-6·Hyaluronan Complex by Modeling and NMR Spectroscopy: Insights into other members of the link module superfamily. Journal of Biological Chemistry, 280(18), 18189–18201. https://doi.org/10.1074/jbc.M414343200

Blundell, C. D., DeAngelis, P. L., Day, A. J., & Almond, A. (2004). Use of ^15^N-NMR to resolve molecular details in isotopically-enriched carbohydrates: Sequence-specific observations in hyaluronan oligomers up to decasaccharides. Glycobiology, 14(11), 999–1009. https://doi.org/10.1093/glycob/cwh117

Blundell, C. D., Reed, M. A. C., & Almond, A. (2006). Complete assignment of hyaluronan oligosaccharides up to hexasaccharides. Carbohydrate Research, 341(17), 2803–2815. https://doi.org/10.1016/j.carres.2006.09.023

Buckley, C., Murphy, E. J., Montgomery, T. R., & Major, I. (2022). Hyaluronic Acid: A Review of the Drug Delivery Capabilities of This Naturally Occurring Polysaccharide. Polymers, 14(17), Article 17. https://doi.org/10.3390/polym14173442

Cowman, M. K., Schmidt, T. A., Raghavan, P., & Stecco, A. (2015). Viscoelastic Properties of Hyaluronan in Physiological Conditions. F1000Research, 4, 622. https://doi.org/10.12688/f1000research.6885.1

D’Agostino, A., Stellavato, A., Busico, T., Papa, A., Tirino, V., Papaccio, G., La Gatta, A., De Rosa, M., & Schiraldi, C. (2015). In vitro analysis of the effects on wound healing of high- and low-molecular weight chains of hyaluronan and their hybrid H-HA/L-HA complexes. BMC Cell Biology, 16(1), 19. https://doi.org/10.1186/s12860-015-0064-6

DeAngelis, P. L., Gunay, N. S., Toida, T., Mao, W., & Linhardt, R. J. (2002). Identification of the capsular polysaccharides of Type D and F Pasteurella multocida as unmodified heparin and chondroitin, respectively. Carbohydrate Research, 337(17), 1547–1552. https://doi.org/10.1016/S0008-6215(02)00219-7

Delaglio, F., Grzesiek, S., Vuister, GeertenW., Zhu, G., Pfeifer, J., & Bax, A. (1995). NMRPipe: A multidimensional spectral processing system based on UNIX pipes. Journal of Biomolecular NMR, 6(3). https://doi.org/10.1007/BF00197809

Donati, A., Magnani, A., Bonechi, C., Barbucci, R., & Rossi, C. (2001). Solution structure of hyaluronic acid oligomers by experimental and theoretical NMR, and molecular dynamics simulation. Biopolymers, 59(6), 434–445. https://doi.org/10.1002/1097-0282(200111)59:6<434::AID-BIP1048>3.0.CO;2-4

Fong Chong, B., & Nielsen, L. K. (2003). Aerobic cultivation of Streptococcus zooepidemicus and the role of NADH oxidase. Biochemical Engineering Journal, 16(2), 153–162. https://doi.org/10.1016/S1369-703X(03)00031-7

Fraser, J. R. E., Laurent, T. C., & Laurent, U. B. G. (1997). Hyaluronan: Its nature, distribution, functions and turnover. Journal of Internal Medicine, 242(1), 27–33. https://doi.org/10.1046/j.1365-2796.1997.00170.x

Ghiselli, G. (2017). Drug-Mediated Regulation of Glycosaminoglycan Biosynthesis. Medicinal Research Reviews, 37(5), 1051–1094. https://doi.org/10.1002/med.21429

Harris, R. K., Becker, E. D., De Menezes, S. M. C., Granger, P., Hoffman, R. E., & Zilm, K. W. (2008). Further Conventions for NMR Shielding and Chemical Shifts (IUPAC Recommendations 2008). Magnetic Resonance in Chemistry, 46(6), 582–598. https://doi.org/10.1002/mrc.2225

Jaroniec, C. P., Filip, C., & Griffin, R. G. (2002). 3D TEDOR NMR Experiments for the Simultaneous Measurement of Multiple Carbon−Nitrogen Distances in Uniformly ^13^ C, ^15^ N-Labeled Solids. Journal of the American Chemical Society, 124(36), 10728–10742. https://doi.org/10.1021/ja026385y

Kass, E. H., & Seastone, C. V. (1944). The rôle of the mucoid polysaccharide (hyaluronic acid) in the virulence of group a hemolytic streptococci. Journal of Experimental Medicine, 79(3), 319–330. https://doi.org/10.1084/jem.79.3.319

Kirui, A., Zhao, W., Deligey, F., Yang, H., Kang, X., Mentink-Vigier, F., & Wang, T. (2022). Carbohydrate-aromatic interface and molecular architecture of lignocellulose. Nature Communications, 13(1), Article 1. https://doi.org/10.1038/s41467-022-28165-3

Liu, L., Liu, Y., Li, J., Du, G., & Chen, J. (2011). Microbial production of hyaluronic acid: Current state, challenges, and perspectives. Microbial Cell Factories, 10(1), 99. https://doi.org/10.1186/1475-2859-10-99

Lu, J. F., Zhu, Y., Sun, H. L., Liang, S., Leng, F. F., & Li, H. Y. (2016). Highly efficient production of hyaluronic acid by Streptococcus zooepidemicus R42 derived from heterologous expression of bacterial haemoglobin and mutant selection. Letters in Applied Microbiology, 62(4), 316–322. https://doi.org/10.1111/lam.12546

Mahoney, D. J., Aplin, R. T., Calabro, A., Hascall, V. C., & Day, A. J. (2001). Novel methods for the preparation and characterization of hyaluronan oligosaccharides of defined length. Glycobiology, 11(12), 1025–1033. https://doi.org/10.1093/glycob/11.12.1025

Mandal, A., Boatz, J. C., Wheeler, T., & van der Wel, P. C. A. (2017). On the use of ultracentrifugal devices for routine sample preparation in biomolecular magic-angle-spinning NMR. Journal of Biomolecular NMR, 67(3), 165–178. https://doi.org/10.1007/s10858-017-0089-6

Marei, W. F., Ghafari, F., & Fouladi-Nashta, A. A. (2012). Role of hyaluronic acid in maturation and further early embryo development of bovine oocytes. Theriogenology, 78(3), 670– 677. https://doi.org/10.1016/j.theriogenology.2012.03.013

Matlahov, I., & van der Wel, P. C. A. (2018). Hidden motions and motion-induced invisibility: Dynamics-based spectral editing in solid-state NMR. Methods (San Diego, Calif.), 148, 123–135. https://doi.org/10.1016/j.ymeth.2018.04.015

Mausolf, A., Jungmann, J., Robenek, H., & Prehm, P. (1990). Shedding of hyaluronate synthase from streptococci. Biochemical Journal, 267(1), 191–196. https://www.ncbi.nlm.nih.gov/pmc/articles/PMC1131263/

Murgoci, A., & Duer, M. (2021). Molecular conformations and dynamics in the extracellular matrix of mammalian structural tissues: Solid-state NMR spectroscopy approaches. Matrix Biology Plus, 12, 100086. https://doi.org/10.1016/j.mbplus.2021.100086

Nestor, G., & Sandström, C. (2017). NMR study of hydroxy and amide protons in hyaluronan polymers. Carbohydrate Polymers, 157, 920–928. https://doi.org/10.1016/j.carbpol.2016.10.005

Patching, S. (2016). NMR-Active Nuclei for Biological and Biomedical Applications. Journal of Diagnostic Imaging in Therapy, 3(1), 7–48. https://doi.org/10.17229/jdit.2016-0618-021

Patil, K. P., Kamalja, K. K., & Chaudhari, B. L. (2011). Optimization of medium components for hyaluronic acid production by Streptococcus zooepidemicus MTCC 3523 using a statistical approach. Carbohydrate Polymers, 86(4), 1573–1577. https://doi.org/10.1016/j.carbpol.2011.06.065

Pérez García, M., Wang, T., Salazar, A., Zabotina, O. A., & Hong, M. (2012). Multidimensional solid-state NMR studies of the structure and dynamics of pectic polysaccharides in uniformly 13C-labeled Arabidopsis primary cell walls. Magnetic Resonance in Chemistry, 50(8), 539–550. https://doi.org/10.1002/mrc.3836

Pomin, V. H., Sharp, J. S., Li, X., Wang, L., & Prestegard, J. H. (2010). Characterization of Glycosaminoglycans by ^15^ N NMR Spectroscopy and in Vivo Isotopic Labeling. Analytical Chemistry, 82(10), 4078–4088. https://doi.org/10.1021/ac1001383

Pouyani, T., Harbison, G. S., & Prestwich, G. D. (1994). Novel Hydrogels of Hyaluronic Acid: Synthesis, Surface Morphology, and Solid-State NMR. Journal of the American Chemical Society, 116(17), 7515–7522. https://doi.org/10.1021/ja00096a007

Rivera Starr, C., & Engleberg, N. C. (2006). Role of Hyaluronidase in Subcutaneous Spread and Growth of Group A Streptococcus. Infection and Immunity, 74(1), 40–48. https://doi.org/10.1128/IAI.74.1.40-48.2006

Samadi, M., Khodabandeh Shahraky, M., Tabandeh, F., Aminzadeh, S., & Dina, M. (2022). Enhanced hyaluronic acid production in Streptococcus zooepidemicus by an optimized culture medium containing hyaluronidase inhibitor. Preparative Biochemistry & Biotechnology, 52(4), 413–423. https://doi.org/10.1080/10826068.2021.1955710

Scholz, I., Hodgkinson, P., Meier, B. H., & Ernst, M. (2009). Understanding two-pulse phase-modulated decoupling in solid-state NMR. The Journal of Chemical Physics, 130(11), 114510. https://doi.org/10.1063/1.3086936

Scott, J. E., & Heatley, F. (1999). Hyaluronan forms specific stable tertiary structures in aqueous solution: A ^13^C NMR study. Proceedings of the National Academy of Sciences, 96(9), 4850–4855. https://doi.org/10.1073/pnas.96.9.4850

Snetkov, P., Zakharova, K., Morozkina, S., Olekhnovich, R., & Uspenskaya, M. (2020). Hyaluronic Acid: The Influence of Molecular Weight on Structural, Physical, Physico-Chemical, and Degradable Properties of Biopolymer. Polymers, 12(8), 1800. https://doi.org/10.3390/polym12081800

Stevens, T. J., Fogh, R. H., Boucher, W., Higman, V. A., Eisenmenger, F., Bardiaux, B., van Rossum, B.-J., Oschkinat, H., & Laue, E. D. (2011). A software framework for analysing solid-state MAS NMR data. Journal of Biomolecular NMR, 51(4), 437–447. https://doi.org/10.1007/s10858-011-9569-2

Takasugi, M., Firsanov, D., Tombline, G., Ning, H., Ablaeva, J., Seluanov, A., & Gorbunova, V. (2020). Naked mole-rat very-high-molecular-mass hyaluronan exhibits superior cytoprotective properties. Nature Communications, 11(1), Article 1. https://doi.org/10.1038/s41467-020-16050-w

Takegoshi, K., Nakamura, S., & Terao, T. (2001). ^13^C–^1^H dipolar-assisted rotational resonance in magic-angle spinning NMR. Chemical Physics Letters, 344(5), 631–637. https://doi.org/10.1016/S0009-2614(01)00791-6

Tang, H., Belton, P. S., Ng, A., & Ryden, P. (1999). ^13^C MAS NMR Studies of the Effects of Hydration on the Cell Walls of Potatoes and Chinese Water Chestnuts. Journal of Agricultural and Food Chemistry, 47(2), 510–517. https://doi.org/10.1021/jf980553h

Taweechat, P., Pandey, R. B., & Sompornpisut, P. (2020). Conformation, flexibility and hydration of hyaluronic acid by molecular dynamics simulations. Carbohydrate Research, 493, 108026. https://doi.org/10.1016/j.carres.2020.108026

Thomas, F., Duff, N. L., Leroux, C., Dartevelle, L., & Riera, P. (2020). Chapter One—Isotopic labeling of cultured macroalgae and isolation of ^13^C-labeled cell wall polysaccharides for trophic investigations. In N. Bourgougnon (Ed.), Advances in Botanical Research (Vol. 95, pp. 1–17). Academic Press. https://doi.org/10.1016/bs.abr.2019.11.005

Ucm, R., Aem, M., Lhb, Z., Kumar, V., Taherzadeh, M. J., Garlapati, V. K., & Chandel, A. K. (2022). Comprehensive review on biotechnological production of hyaluronic acid: Status, innovation, market and applications. Bioengineered, 13(4), 9645–9661. https://doi.org/10.1080/21655979.2022.2057760

Wessels, M. R., Moses, A. E., Goldberg, J. B., & DiCesare, T. J. (1991). Hyaluronic acid capsule is a virulence factor for mucoid group A streptococci. Proceedings of the National Academy of Sciences of the United States of America, 88(19), 8317–8321. https://www.ncbi.nlm.nih.gov/pmc/articles/PMC52499/

Wu, C.-S., & Liao, H.-T. (2005). A new biodegradable blends prepared from polylactide and hyaluronic acid. Polymer, 46(23), 10017–10026. https://doi.org/10.1016/j.polymer.2005.08.056

Xu, H., Ito, T., Tawada, A., Maeda, H., Yamanokuchi, H., Isahara, K., Yoshida, K., Uchiyama, Y., & Asari, A. (2002). Effect of hyaluronan oligosaccharides on the expression of heat shock protein 72. The Journal of Biological Chemistry, 277(19), 17308–17314. https://doi.org/10.1074/jbc.M112371200

