## Supplemental Information file for "Production of isotopically enriched high molecular weight hyaluronic acid and characterization by solid-state NMR"

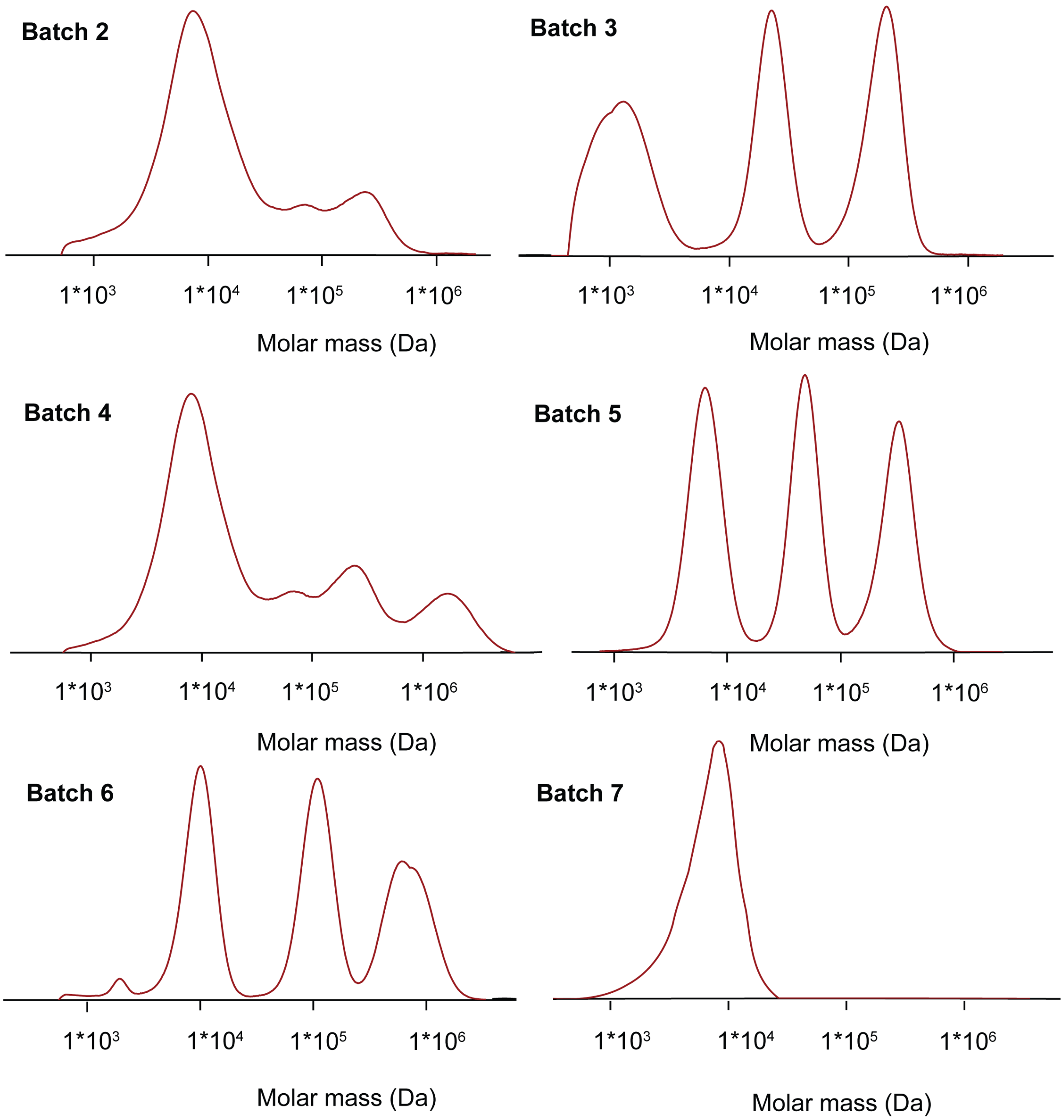


**Figure S1:** GPC molecular mass distribution of different batches of HA. One peak at 10 kDa and three more peaks at higher MW values (in the range of 100 kDa to 1.5 MDa) can be identified. Batch 2, 3, 4, 5, 6 and 7 from Table 1 are reported. Note that the intensity (y axis here omitted) cannot be considered as a reliable indicator of the relative amount of material corresponding to a certain MW.

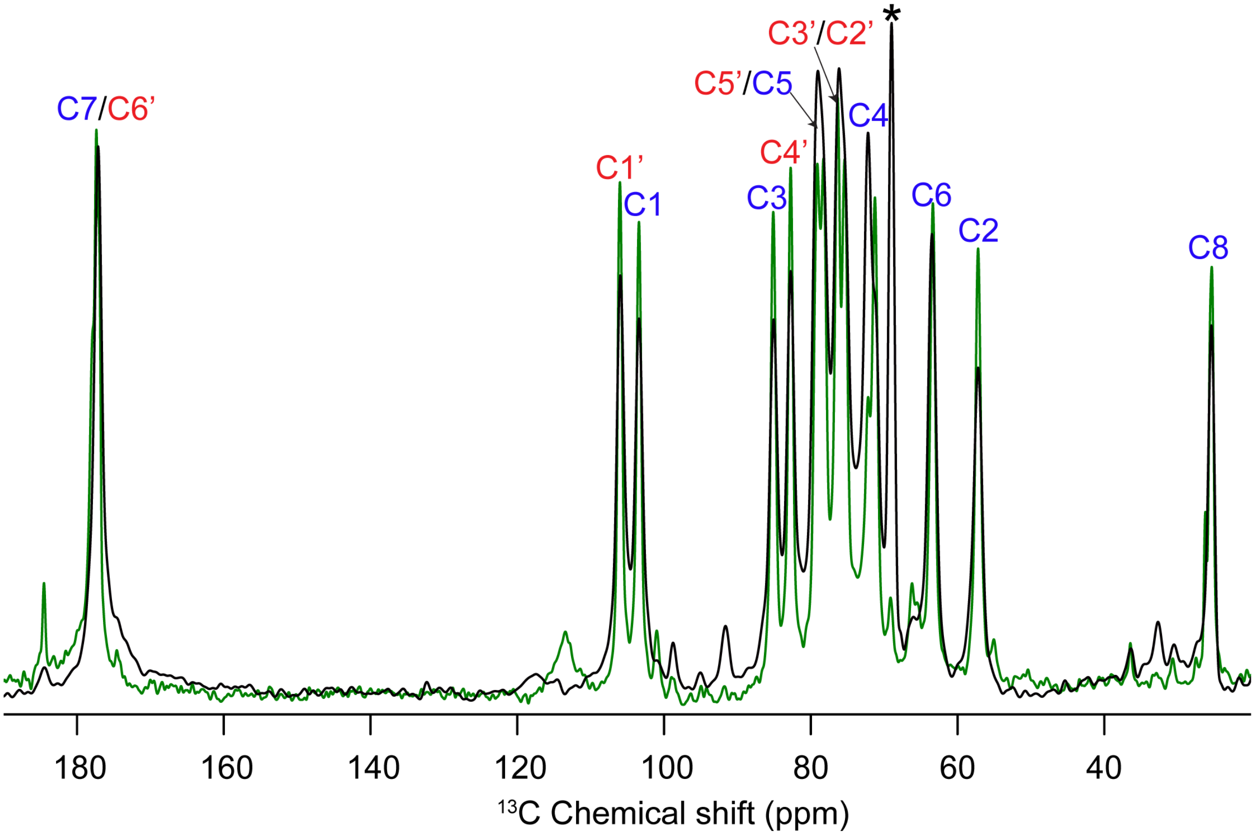


**Figure S2:** Overlaid 1D $^{13}$C DE MAS NMR spectra of $^{13}$C-labelled HMW HA with and without contaminant, from batches 7 and 4 in Table 1. The peak labeled with an asterisk (*) is attributed to the contaminant.

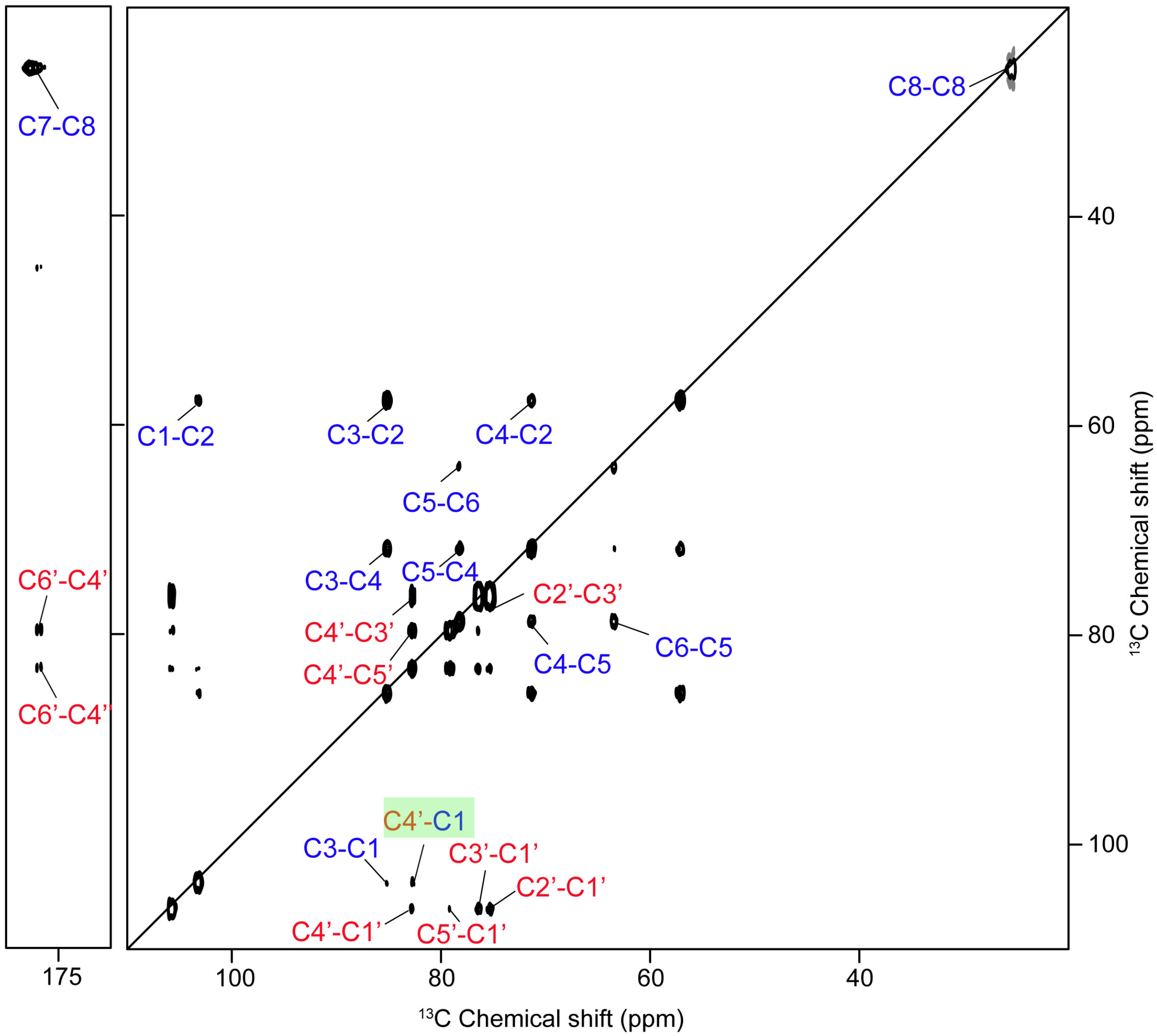


**Figure S3:** 2D $^{13}C$-$^{13}C$ DARR ssNMR spectrum of $^{13}C$ HMWHA with resonance assignment (see also Figure 5 in the main text). The red labels are for the GlcA and blue labels are for the GlcNAc moieties of the HA polysaccharide. The green shaded rectangular box highlights a cross peak between the two moieties (1-4 glycosidic linkage). Cross-peaks involving carbonyl groups are visible in the left-most figure panel.

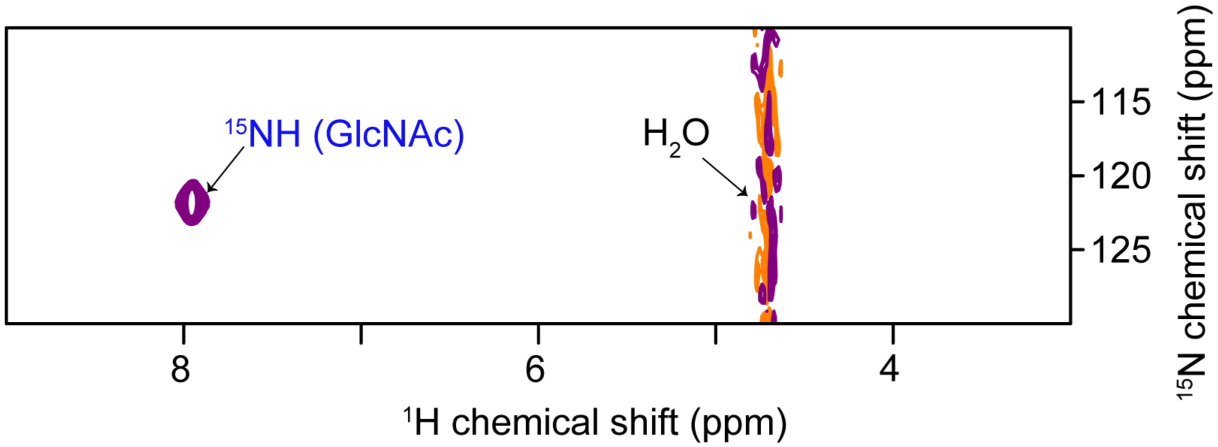


**Figure S4:** 2D $^{1}H$-$^{15}N$ HSQC (Heteronuclear Single Quantum Coherence) spectrum of $^{15}N$-$^{13}C$ labelled HA. The amide $^{1}H$ and $^{15}N$ chemical shifts are indicated as $^{15}NH$.

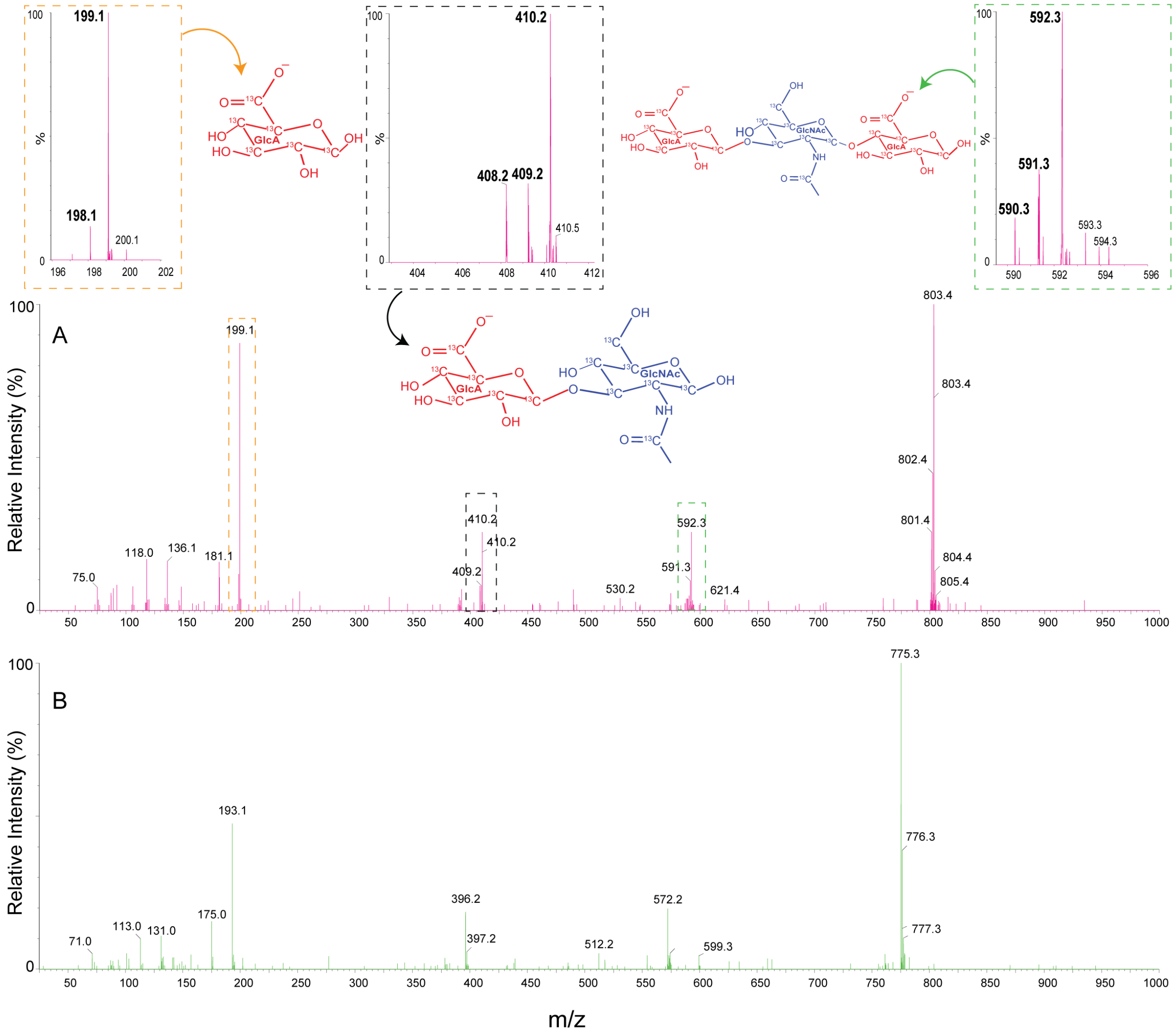


**Figure S5:** Negative ion LC-ESI-MS/MS Fragmentation spectra of $^{13}C$ isotopically labelled (batch 6) and unlabeled HA (batch 1). (A) MS/MS fragmentation spectrum of $^{13}C$ labelled HA (pink) and (B), unlabeled HA (green). Spectral insets in the dashed rectangular boxes are the zoomed-in regions of three main fragments indicating the mass of 199, 410, and 592 Da which correspond to one (GlcA), two (GlcA-GlcNAc) and three moieties (GlcA-GlcNAc-GlcA) of HA. The percentage of partial incorporation was analyzed using the mass of $^{13}C$-lablled GlcA fragment (199.1, 198.1).

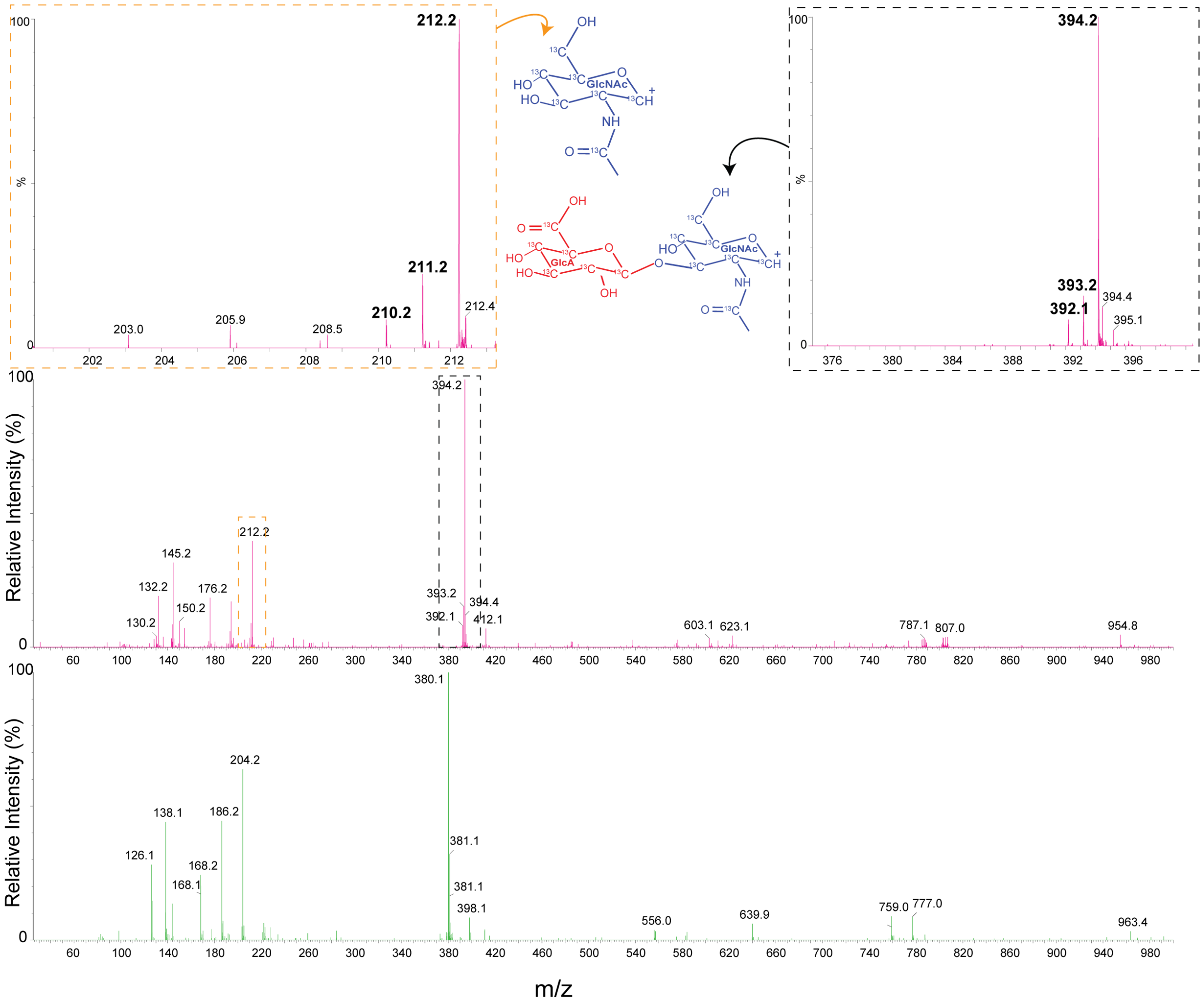


**Figure S6:** Positive mode LC-ESI-MS/MS Fragmentation spectra of $^{13}C$ isotopically labelled (batch 6) and unlabeled HA (batch 1). (A) MS/MS fragmentation spectrum of $^{13}C$ labelled HA (pink) and (B), unlabeled HA (green). Spectral insets in the dashed rectangular boxes are the zoomed-in rgions of two main fragments indicating the mass of 212.2 and 394.2 which correspond to one (GlcNAc) and two moieties (GlcA-GlcNAc) of HA. The percentage of partial incorporation was analyzed using the mass of $^{13}C$-labelled GlcNAc fragment (212.2, 211.2).

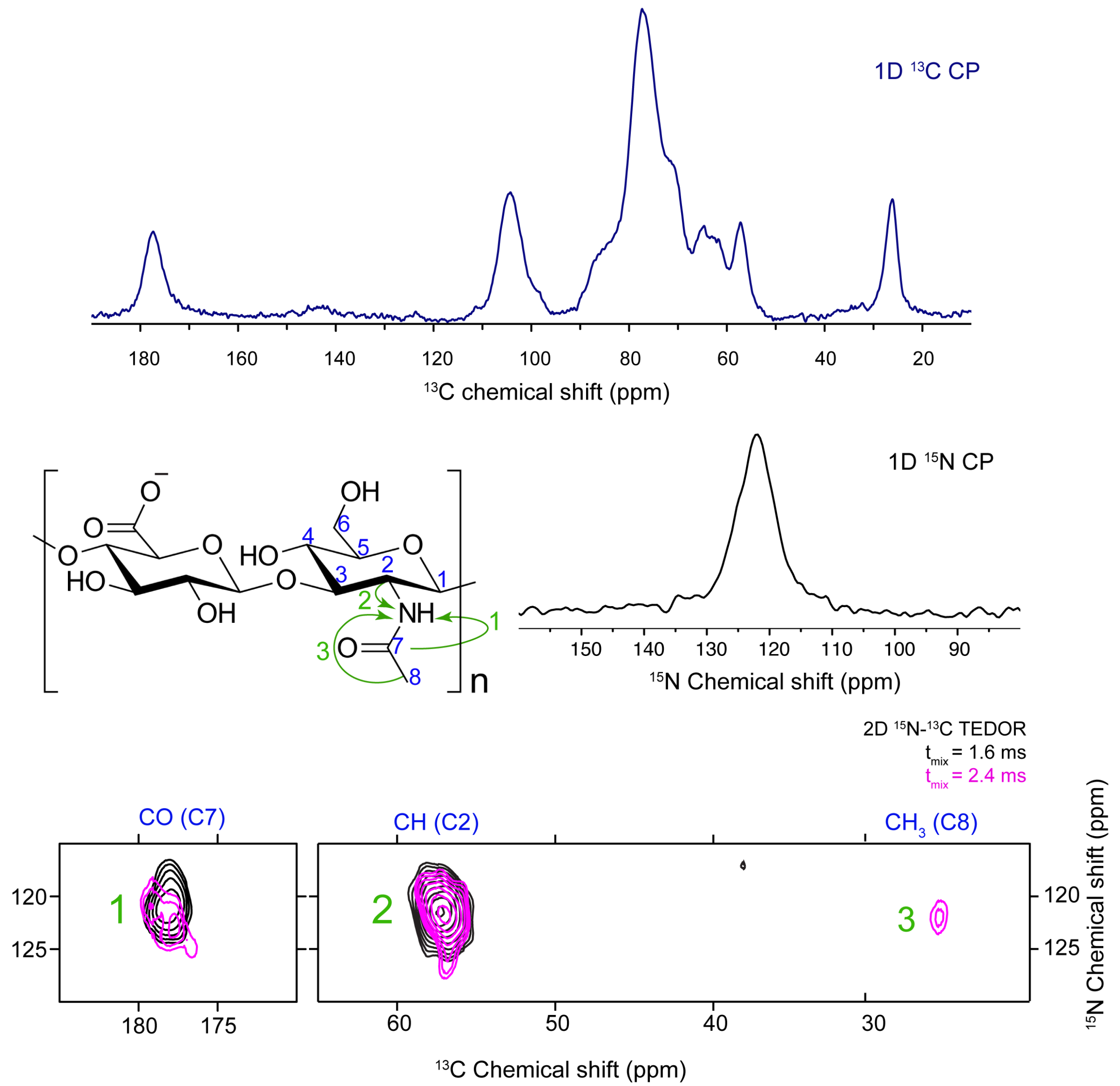


**Figure S7.** 1D and 2D MAS $^{13}C$, $^{15}N$ and $^{15}N$-$^{13}C$ NMR spectra of $^{15}N$,$^{13}C$ HMW HA (batch 8, table 1). Top (blue) and the middle right panel (black) are 1D $^{13}C$ and 1D $^{15}N$ CP-MAS NMR spectra of HA. Bottom panel is a zoomed-in regions of 2D TEDOR spectra recorded with two different mixing times. Blue labels (on top of TEDOR spectrum) and green curvy arrows in the HA chemical structure shows correlations of carbons with amide nitrogen.

**Table S1.** Observed molecular masses of unlabeled and isotopically labeled HA tetrasaccharides obtained from enzymatic digestion, and determination of $^{13}C$ and $^{15}N$ incorporation. From the reported values, 97% of $^{13}C$ and 52% of $^{15}N$ incorporation can be calculated. The overall percentage of $^{13}C$ and $^{15}N$ incorporation is calculated as the weighted average among the different species (M, M-1, [M]-2, etc.) of the ratio between the number of total $^{13}C$ and $^{15}N$ atoms incorporated and the number of C and N atoms present in tetrasaccharide unit. The analyzed HA batches were obtained in conditions as from batch 1, 4 and 8 in Table 1.

| Species | Observed mass | Difference (Da) | Isotopic adduct | Relative intensity (%) | Relative abundance (%) | $^{13}C$ / $^{15}N$ atoms |
|---|---|---|---|---|---|---|
| **$HA_4$** | 774.9 | 0 | [M] | 100 | 32.63 | 0 |
| | 775.9 | +1 | [M+H] | 28.39 | 9.26 | 0 |
| | 792.5 | +18 | [M+$H_2O$] | 67.9 | 22.16 | 0 |
| | 796.9 | +22 | [M-H+$Na^+$] | 79.87 | 26.06 | 0 |
| **$^{13}C$-$HA_4$** | 802.9 | 0 | [M] | 100 | 52.35 | 28 |
| | 801.9 | -1 | [M] – 1x $^{13}C$ | 29.6 | 15.49 | 27 |
| | 800.9 | -2 | [M] – 2x $^{13}C$ | 41.5 | 21.72 | 26 |
| | 799.9 | -3 | [M] – 3x $^{13}C$ | 12.4 | 6.49 | 25 |
| | 798.9 | -4 | [M] – 4x $^{13}C$ | 7.54 | 3.95 | 24 |
| **$^{13}C$,$^{15}N$-$HA_4$** | 804.9 | 0 | [M] | 100 | 34.46 | 30 |
| | 803.9 | -1 | [M] – 1x $^{15}N$* | 99 | 34.11 | 29 |
| | 802.9 | -2 | [M] – 2x $^{15}N$* | 41 | 14.13 | 28 |
| | 801.9 | -3 | [M] – (2x $^{15}N$ + 1x $^{13}C$)* | 25.8 | 8.89 | 27 |
| | 800.9 | -4 | [M] – (2x $^{15}N$ + 2x $^{13}C$)* | 9.3 | 3.20 | 26 |
| | 799.9 | -5 | [M] – (2x $^{15}N$ + 3x $^{13}C$)* | 7.6 | 2.62 | 25 |
| | 798.9 | -6 | [M] – (2x $^{15}N$ + 4x $^{13}C$)* | 7.5 | 2.58 | 24 |

M denotes the mass of HA tetrasaccharide. *These adducts are based on the assumption that $^{15}N$ incorporation is less efficient than $^{13}C$ incorporation, having other $^{15}N$ sources available in the media apart from $^{15}N$ ammonium chloride.
